# Isopropoxy benzene guanidine kills *Staphylococcus aureus* without detectable resistance

**DOI:** 10.1101/2020.03.10.986687

**Authors:** Xiufeng Zhang, Xianfeng Peng, Yixing Lu, Jie Hao, Fangping Li, Zonghua Qin, Wenguang Xiong, Zhenling Zeng

**Affiliations:** Guangdong Provincial Key Laboratory of Veterinary Drugs Development and Safety Evaluation, South China Agricultural University, Guangzhou 510642, China; National Risk Assessment Laboratory for Antimicrobial Resistance of Animal Original Bacteria, South China Agricultural University, Guangzhou 510642, China; Guangdong Laboratory for Lingnan Modern Agriculture, Guangzhou, 510642, China; Guangzhou Insighter Biotechnology Co., Ltd, Guangzhou 510642, China; Guangdong Provincial Key Laboratory of Plant Molecular Breeding, State Key Laboratory for Conservation and Utilization of Subtropical Agro-Bioresources, South China Agricultural University, Guangzhou 510642, China

## Abstract

Antibiotic resistance is a serious public health crisis. The challenge caused by *Staphylococcus aureus* infections clearly urges the development of novel antimicrobial therapy. Drug repurposing has emerged as an alternative approach to rapidly identify effective drugs against multidrug-resistant bacteria. Recently, substituted benzene guanidine compounds have been used as leading structures to discover new promising drugs in both synthetic and medicinal chemistry. We investigated the antimicrobial activity of an analog of substituted benzene guanidine compounds (isopropoxy benzene guanidine) and further explored its antibacterial mechanism against *S. aureus*. Isopropoxy benzene guanidine had a MIC of 0.125-4μg/ml against *S. aureus* and displayed potent activity against *S. aureus* by disrupting cell membrane. Unlike conventional antibiotics, repeated use of isopropoxy benzene guanidine had a low probability of resistance section. The most substantial isopropoxy benzene guanidine-induced changes occurred in transcript levels of membrane transport functions-regulated genes, and genes involved in purine- and pyrimidine-synthesis pathway and virulence factors. Furthermore, *in vivo* studies demonstrated that isopropoxy benzene guanidine is capable of treating invasive MRSA infections. These findings provided strong evidence that isopropoxy benzene guanidine represents a new chemical lead for novel antibacterial agent against multidrug-resistant *S. aureus* infections.

## Introduction

Antibiotic resistance is one of the most prominent public health challenges of our time. Adding to this dire situation, the antibiotic pipeline has slowed to a drip over the past 30 years (1). *Staphylococcus aureus*, one member of “ESKAPE” pathogens, is currently the most clinically important multidrug-resistant pathogen and responsible for life-threatening infections (2). Meanwhile, the alarming increase in the prevalence of global spread clones in *S. aureus* resistant to nearly all antibiotics is a major public health concern (3). Hence, there is a dire need to develop novel antimicrobial compounds, as well as are tolerated with low propensity for resistance development (4–5).

However, antimicrobial drug discovery is a costly, complex and time-consuming process. Until June 2019, only about 42 new antibiotics with the potential to treat serious bacterial infections are in clinical development (6). Given historical data, only 1 in 5 infectious disease products are likely to receive regulatory approval during the next 7 years (7). To address this problem, repurposing or repositioning drugs have emerged as an innovation stream of pharmaceutical development and gained great success in treating various infectious diseases (8–10).

The majority of recently approved agents have been developed from existing drug classes, rather than new chemical classes (11). Compounds such as aminoguanidines have tremendous potential for new chemical synthesis and biological applications (12–13). Substituted benzene guanidine compounds, which belong to aminoguanidines, have undergone a mammoth amount of screening and testing to discover new promising drugs in both synthetic and medicinal chemistry (14–15). For example, robenidine synthesized as an anticoccidial agent was widely used to prevent coccidian infections since the early 1970s (16). Recently, robenidine analogues have been shown to inhibit Gram-positive antibacterial agents, including *S. aureus* and vancomycin-resistant *Enterococci* (17). Moreover, robenidine analog NCL195 could disrupt the cell membrane potential in *Streptococcus pneumonia* and *Staphylococcus aureus* (18). Hence, the design of substituted benzene guanidine compounds as anti-Gram-positive bacteria agents is highly desirable.

In our previous study, we screened a compound from substituted benzene guanidine derivative, isopropoxy benzene guanidine (IBG), as anti-*Enterococci* agent by disrupting their cell membrane potential (19). We further found it also had potent antibacterial activity against *S. aureus*. However, there is no in-depth evaluation of this compound against *S. aureus.*

In this study, we characterized the response of *S. aureus* ATCC 29213 treated with IBG using phenotypic assays and transcriptomics. The *in vivo* treatment efficacy of IBG was also investigated in a mouse septicemia model.

## Materials and methods

### Bacterial strains, growth conditions

*S. aureus* strain ATCC 29213, methicillin-resistant *S. aureus* (MRSA) ATCC 43300, 105 clinical *S. aureus* isolates (including 55 MRSA strains (20)) were used in this study. All strains were grown in Mueller-Hinton (MH) broth. Multilocus sequence typing (MLST) was conducted according to the reference MLST database.

### Antimicrobial agents and chemicals

Vancomycin, gentamicin, ciprofloxacin, linezolid, amikacin, trixon X-100 and *tris*-(hydroxrymethyl) aminomethane (Tris) were purchased from Sangon Biotech (Shanghai, China). Isopropoxy benzene guanidine (IBG) (batch number 20150506, content 99.9%) was synthesized by Guangzhou Insighter Biotechnology (Guangzhou, China). Fetal bovine serum (FBS) was from Zhejiang Tianhang Biotechnology (Zhejing, China). Dimethyl Sulphoxide (DMSO) (Dmreagent, Tianjing, China) was utilized as solvent to dissolve IBG. Anti-infective detergent benzalkonium chloride (BAC) was purchased from Aladdin Industrial Corporation (Shanghai, China). N,N’-dicyclohexylcarbodiimide (DCCD) was from Macklin Biochemical (Shanghai, China). SYTOX^®^ Green Nucleic Acid Stain and 3, ‘-diethyloxacarbocyanine iodide (DiOC_2_ (3)) (Thermo Fisher Scientific, Germany) were used as Molecular Probes.

### Minimum inhibitory concentration (MIC) assay

The MIC of IBG against *S. aureus* strain ATCC 29213, MRSA ATCC 43300, 105 clinical *S. aureus* was determined by broth microdilution according to CLSI guidelines (21). Cell concentration was adjusted to approximately 5×10^5^ CFU/ml. After 16-20 h of incubation at 37°C, the MIC was defined as the lowest concentration of antibiotic with no visible growth. The presence of FBS on IBG activity against *S. aureus* was also tested. Experiments were performed with biological replicates.

### Antibacterial activity with detergents or ATPase inhibitors

The antibacterial activity of IBG in the presence and absence of detergents (Triton X-100 and Tris) and ATPase-inhibiting agent (DCCD) were investigated. Triton X-100 and Tris could increase permeability of outer membrane, DCCD can decrease bacterial ATP levels by disrupting electrochemical proton gradients. The antibacterial activity of IBG against *S. aureus* strain ATCC 29213 and MRSA ATCC 43300 was measured in the presence of 0.001% Triton X-100, 125μg/ml Tris or 250 μg/ml DCCD. After incubating for 18h, the OD_600nm_ was measured using a multifunctional microplate reader (Thermo Multiskan FC, Molecular Devices, USA). The assays were conducted in triplicates.

### Minimum bactericidal concentration (MBC)

The MBC was further determined recommended by CLSI. Briefly, 20 μl aliquots were taken from all the wells from MIC microbroth plate that had been incubated for 20h and then spotted onto MH agar. The colonies enumerated after incubating for 24 h at 37°C. The MBC is defined as the lowest concentration where a 99.9% colony count reduction was observed (22). Experiments were performed in triplicates.

### Killing kinetics assay

The time-dependent killing for *S. aureus* ATCC 29213 and MRSA 43300 with IBG, vancomycin (VAN) were investigated previously (23). An overnight culture of cells was diluted 110000 in MH broth and incubated at 37°C, 180 rpm for 4 h until the cell suspension was about 1×10^7^CFU/ml and challenged with IBG, VAN at 10× MIC at 37°C and 225 r.p.m. At different time points, 100 μl aliquots were ten-fold serially diluted and plot-plated with MH agar plates. After incubating the plates at 37°C overnight, the plates with around 30 to 300 colonies were counted and CFU/ml for each time point was calculated. To determine bacterial lysis, 5ml of exponential-phase *S. aureus* ATCC 29213 culture (OD_600nm_ ~0.4) was treated with 10× MIC of IBG, VAN and BAC (positive control for bacterial lysis) for 4 h (24). Every hour, 200μl of each sample was added to 96-well microtiter plate and the OD_600nm_ was measured using a multifunctional microplate reader. All the experiments were replicated.

### Resistance selection

In order to select IBG-resistant mutants, ~10^10^ CFU of *S. aureus* ATCC 29213 cells were plated onto MH agar containing 2.5×, 5×, and 10×MIC of IBG. After 48 h of incubation at 37°C, resistant colonies were calculated and the MICs of IBG were determined. When this approach proved unsuccessful, development of resistant mutants by serial passage in liquid medium was conducted previously (23). *S. aureus* ATCC 29213 at exponential phase were diluted to OD_600_ of 0.01 in 2 ml of MHB. Cells were treated with IBG at 0.25×MIC, 0.5×MIC, 1×MIC, 2×MIC and 4×MIC. At 24 hour intervals, the cultures from the second highest concentrations that allowed bacteria growth were diluted 1100 into fresh MH broth containing 0.25×MIC, 0.5×MIC, 1×MIC, 2×MIC and 4×MIC of IBG. The rest of the culture was stored in 30% glycerol at −80°C. Serial passage was performed on three independent cultures (SP1, SP2, SP3) for 100 days, and a separate ciprofloxacin selection served as a control. Any cultures that grew at higher than the MIC levels were passaged on drug free MHA plates and the MIC was then determined by broth microdilution.

### Genomic DNA extraction, library preparation and genome sequencing

*S. aureus* ATCC 29213 genomic DNAs from days 0, 58, 99 from SP1, days 58, 100 from SP2 and days 34, 66, 100 from SP3 were extracted using a Hipure bacterial DNA kit (Magen, Shanghai, China). Genomic DNA was quantified using NanoDrop 2000 spectrophotometer (Thermo Fisher Scientific Inc.) and agarose gel electrophoresis. A paired-end sequencing library (2×250bp) was created using a VAHTS Universal DNA Library Prep kit for Illumina^®^ (Illumina, San Diego, CA, USA) and sequenced within an Illumina HiSeq system (Illumina Inc.).

### Genome assembly, annotation and variant calling

All the genomes were *de novo* assembled using CLC Genomics Workbench 10.1 (CLC Bio, Aarhus, Denmark). All the genomes were annotated by using Prokka pipeline and then these genomes were subjected to SNP analysis by utilizing snippy pipeline. *S. aureus* ATCC 29213 genome of the starting strain was used as a reference genome in SNP analysis.

### Membrane potential assay

To examine the perturbation of the cell membrane of *S. aureus* by IBG, the membrane potential of the cells was measured by fluorescence spectrometry using fluorescent membrane potential probe DiOC_2_ (3), as described previously (25). *S. aureus* ATCC 29213 cells were grown in LB broth for 12h and resuspended in PBS to OD_600 nm_=0.5. 2 ml of suspension was added in a quartz cuvette and stirred gently for 5 min (with or without addition of 10×MIC, 4×MIC, 1×MIC IBG, 10×MIC ampicillin as control). The fluorescence was measured by a Hitachi F-7000 Fluorescence Spectrometer set at Ex.486 nm/Em. 620 nm, with excitation and emission slit widths at 5 nm and 10 nm, respectively). The background fluorescence of each suspension was monitored about 50s until it plateaued after adding a final concentration of 10 μM DiOC_2_ (3). Cells were re-energized establish a proton motive force with 0.5% glucose and membrane potential was disrupted with the proton ionophore CCCP (10 μM). All assays were performed at least twice.

### Membrane permeability assay

To confirm the integrity of the bacterial membranes, we performed an assay based on the uptake of the fluorescent dye with SYTOX Green (24). Black, clear-bottom, 96-well plates were filled with 50 μl PBS/well containing different concentration of IBG. Exponential-phase *S. aureus* ATCC 29213 cells were washed 3 times with PBS and adjusted to OD_600_ = 0.4 in PBS. SYTOX Green was added to 10 ml bacterial suspension to a final concentration of 5 μM and incubated for 30 min at room temperature in the dark. 50 μl of the bacteria-SYTOX Green mixture was added to the 96-well plates containing IBG and fluorescence was measured using a multifunctional microplate reader, with excitation and emission wavelengths of 485 nm and 525 nm, respectively. All experiments were conducted in duplicate.

### Transmission electron microscopy

Morphological appearance and morphometric analysis of the cell membrane of *S. aureus* ATCC 29213 after exposure to 10×MIC IBG was determined using TEM as described previously (26). Briefly, exponential phase of *S. aureus* ATCC 29213 cells were treated with 10×MIC of IBG or 0.1% DMSO (control) at 37°C for 4h. 1 ml of cells was fixed with 5% glutaraldehyde, incubated for at least 2 h at room temperature and then stored at 4°C. Fixed cells were washed in 0.1 M cacodylate buffer and post-fixed with 1% Osmium tetroxide/1.5% Potassium ferrocyanide for 1 h. Cells were then washed twice in water and subsequently dehydrated in an alcohol gradient series (50%, 70%, 90%, 10 min each 100% 2×10min). Then cells were put in propylene oxide for 1 h and infiltrated overnight in a 11 mixture of propylene oxide and Spurr’s low viscosity resin (Electron Microscopy sciences, Hatfield, PA). The cells were embedded in Spurr’s resin and polymerized at 60°C for 48 h. Ultrathin sections (about 60 nm) were cut on a Reichert Ultracut-S microtome (Leica Microsystem, Wetzlar, Germany), picked up onto copper grids, and stained with lead citrate. Micrographs of the cells were taken using a JEOL 1200EX transmission electron microscope (Harvard Medical School EM facility).

### RNA-Seq transcriptomics

*S. aureus* ATCC 29213 cells were grown to an OD_600_ of 0.4 and treated with 10×MIC IBG for 4h. Samples were collected and preserved with RNA protect (Qiagen, United States). Total RNA of each sample was extracted using TRIzol Reagent (Invitrogen)/RNeasy Mini Kit (Qiagen). Control samples were collected from an antibiotic-free culture. RNA sequencing was conducted by High-Throughput Sequencing Facility at the GENWIZ lnc (Jiangsu, China).

### Bioinformatic analysis

Raw sequence data were underwent quality control using Cutadapt (V1.9.1) and FastQC (V0.10.1). Clean data were aligned with the reference genome of *S. aureus* NCTC 8325 (NCBI accession number NC_007795.1) via software Bowtie2 (v2.1.0) and the gene expression level were estimated by HTSeq (v0.6.1p1). The calculation of fragment per kilobase of exon per million fragments mapped (FPKM) for all genes were performed through Cufflinks (v2.2.1) software. Differential expression genes (DEGs) were screened out by using the DESeq Bioconductor package and defined as those with a change in expression of >2-fold and a corresponding false discovery rate (FDR) of < 0.05. The gene ontology terms and functional pathways were annotated via Gene Ontology (GO) and Kyoto Encyclopedia of Genes and Genome (KEGG), respectively. Cell-PLoc 2.0 was used to analyze the subcellular localization of DEGs (27). ClueGO (2.5.5), a plugin for Cytoscape (3.7.1), was used to visually demonstrate DEGs distribution in KEGG pathway (28).

### Antibiotic synergy test

The checkerboard method was used for determining synergy of IBG with conventional antibiotics (29). Briefly, 2-fold serial dilutions of each IBG were combined with 2-fold serial dilutions of each conventional antibiotic, creating an 10×8 matrix in a 96-well microtiter plate. The fractional inhibitory concentration index (FICI) was calculated as follows FICI = MIC of compound A in combination / MIC of compound A alone + MIC of compound B in combination / MIC of compound B alone. The interaction between two compounds was defined as synergy if FICI<0.5, addition if 0.5≤FICI≤1, no interaction if 1<FICI≤4, antagonism if FICI>4 (30).

### Animal studies

All animal studies were carried out at Animal laboratory center of South China Agricultural University and conformed to institutional animal care and use policies. All animal studies were performed with specific-pathogen-free female KM mice (Southern Medical University, Guangdong, China), 6-8-weeks old, weighing 20±2g.

### Mouse sepsis protection model

IBG was tested against MRSA YXMC004P in a mouse septicemia protection assay to assess its *in vivo* treatment efficacy (31–32). KM female mice were treated intraperitoneally with a dose of 40 mg/kg IBG. After 24h, KM female mice were infected with 0.1ml of bacterial suspension via tail vein injection, a concentration that achieves about 90% mortality within 18 days after infection. At 0.5h and 24 post infection, mice (14 per group) were treated with 40 mg/kg IBG. Infection control mice were intraperitoneally with vancomycin at a clinical dose of 15mg/kg or PBS alone. Mice were monitored for 18 days after infection and the probability determined by non-parametric log-rank test.

### Ethics statement

Mouse studies were approved by the Animal Research Committee of South China Agricultural University (ID (2018)030). All experiments were conducted in full compliance with the guidelines of Guangdong Laboratory Animal Welfare and Ethics and the Institutional Animal Care and Use Committee of the South China Agricultural University.

## Results

### Antimictobial activity of IBG against multidrug-resistant *S. aureus*

IBG exhibited potent activity against all tested *S. aureus* with the MIC range of 0.125-4μg/ml. However, in the presence of 10% FBS, the MICs of IBG against *S. aureus* ATCC 29213 and MRSA 43300 were 8-fold of their MICs (Table 1). When treated with membrane permeability Trixon X100, Tris or ATPase inhibitors DCCD, the OD600 of the suspensions significantly diminished compared to the group treated with IBG alone (Fig. 1).

**Table 1.** Activity of IBG against *Staphylococcus aureus*

| Organism | IBG MIC (µg/ml) | VAN MIC (µg/ml) |
| --- | --- | --- |
| *S. aureus* ATCC 29213 (MSSA) | 4 | 1 |
| *S. Aureus* 29213 + 10% FBS | 32 | 1 |
| MRSA ATCC 43300 | 4 | 1 |
| MRSA 43300 + 10% FBS | 32 | 1 |
| Clinical MSSA (50) | 0.125-4 | 0.5-1 |
| Clinical MRSA (55) | 2-4 | 0.5-1 |

IBG, isopropoxy benzene guanidine; VAN, vancomycin; FBS, fetal bovine serum; MIC, Minimum inhibitory concentration; MSSA, methicillin-sensitive *S. aureus*; MRSA, methicillin-resistant *S.aureus*.

**Fig. 1.**
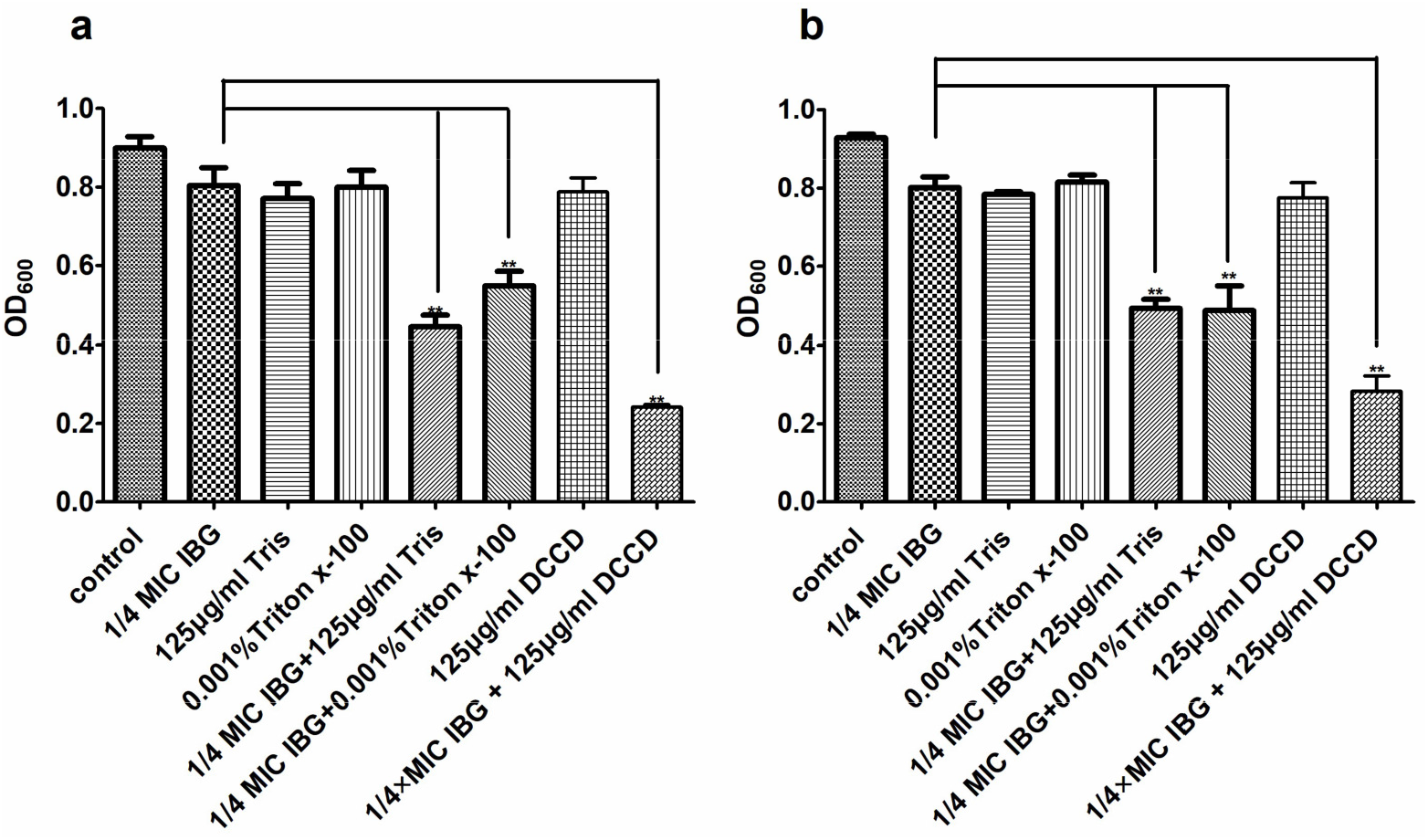
Effects of the membrane-permeabilizing agents (Triton X-100 and Tris) and ATPase-inhibitors (DCCD) on the susceptibility of *S. aureus* to IBG. a, Viability of *S. aureus* ATCC 29213 treated with IBG with Triton X-100, Tris or DCCD. b, Viability of MRSA ATCC 43300 treated with IBG with Triton X-100, Tris or DCCD. IBG, isopropoxy benzene guanidine. ** indicates p<0.01.

### IBG killing kinetics against *S. aureus*

The MBCs of IBG was determined against clinical isolates harbouring different MLST types. Our results showed that the MBC range for IBG against *S. aureus* was 4-8-fold of their MICs (Table 2). Killing kinetics of IBG against *S. aureus* ATCC 29213 and MRSA ATCC 43300 were performed. For *S. aureus* ATCC 29213, we found that IBG and VAN caused a 3.6 and 3.0 log10 cfu/ml decrease in bacterial concentration within 4h, respectively. For MRSA 43300, IBG and VAN reduced a 2.4 and 2.7 log_10_ cfu/ml in the bacterial concentration within 4h, respectively (Fig. 2a, 2b). After 24h, some *S. aureus* regrowth were observed for IBG. We subjected the colonies that grew after 24h for MIC testing we found all colonies returned with 1× MIC for IBG. It was note that the OD600 of the suspensions did not change, which indicated IBG could not cause *S. aureus* lysis (Fig. 2c).

**Table 2.** MBC of IBG against *Staphylococcus aureus*

| Organism | MLST | Spa type | MIC (µg/ml) | MBC (µg/ml) |
| --- | --- | --- | --- | --- |
| MSSA ATCC 29213 |  |  | 4 | 16 |
| MRSA ATCC 43300 |  |  | 4 | 16 |
| MSSA GDM4P080P | 9 | 899 | 4 | 32 |
| MSSA GDE5P004P | 9 | 1775 | 2 | 16 |
| MSSA 131372 | 22 | - | 4 | 16 |
| MSSA 131373 | 7 | - | 4 | 16 |
| MSSA 131374 | 188 | - | 4 | 16 |
| MSSA 131390 | 59 | - | 4 | 16 |
| MSSA 131391 | 5 | - | 4 | 16 |
| MSSA 131403 | 944 | - | 4 | 16 |
| MSSA 131440 | 15 | - | 4 | 16 |
| MSSA 131447 | 6 | - | 0.125 | 1 |
| MSSA 131448 | 88 | - | 4 | 16 |
| MSSA 131449 | 188 | - | 4 | 16 |
| MRSA THPS19-2 | 9 | 899 | 4 | 32 |
| MRSA GDT4P196P | 9 | 1939 | 4 | 16 |
| MRSAGDC6P096P | 398 | 34 | 4 | 16 |
| MRSA SHP6P028P | 398 | 571 | 2 | 8 |
| MRSA 131371 | 338 | - | 4 | 16 |
| MRSA 131404 | 45 | - | 4 | 16 |
| MRSA 131405 | 59 | - | 4 | 16 |
| MRSA 131436 | 1 | - | 4 | 16 |

MBC, Minimum bactericidal concentration; IBG, isopropoxy benzene guanidine; -:undetected; MIC, Minimum inhibitory concentration. MSSA, methicillin-sensitive *S. aureus*; MRSA, methicillin-resistant *S. aureus*.

**Fig. 2.**
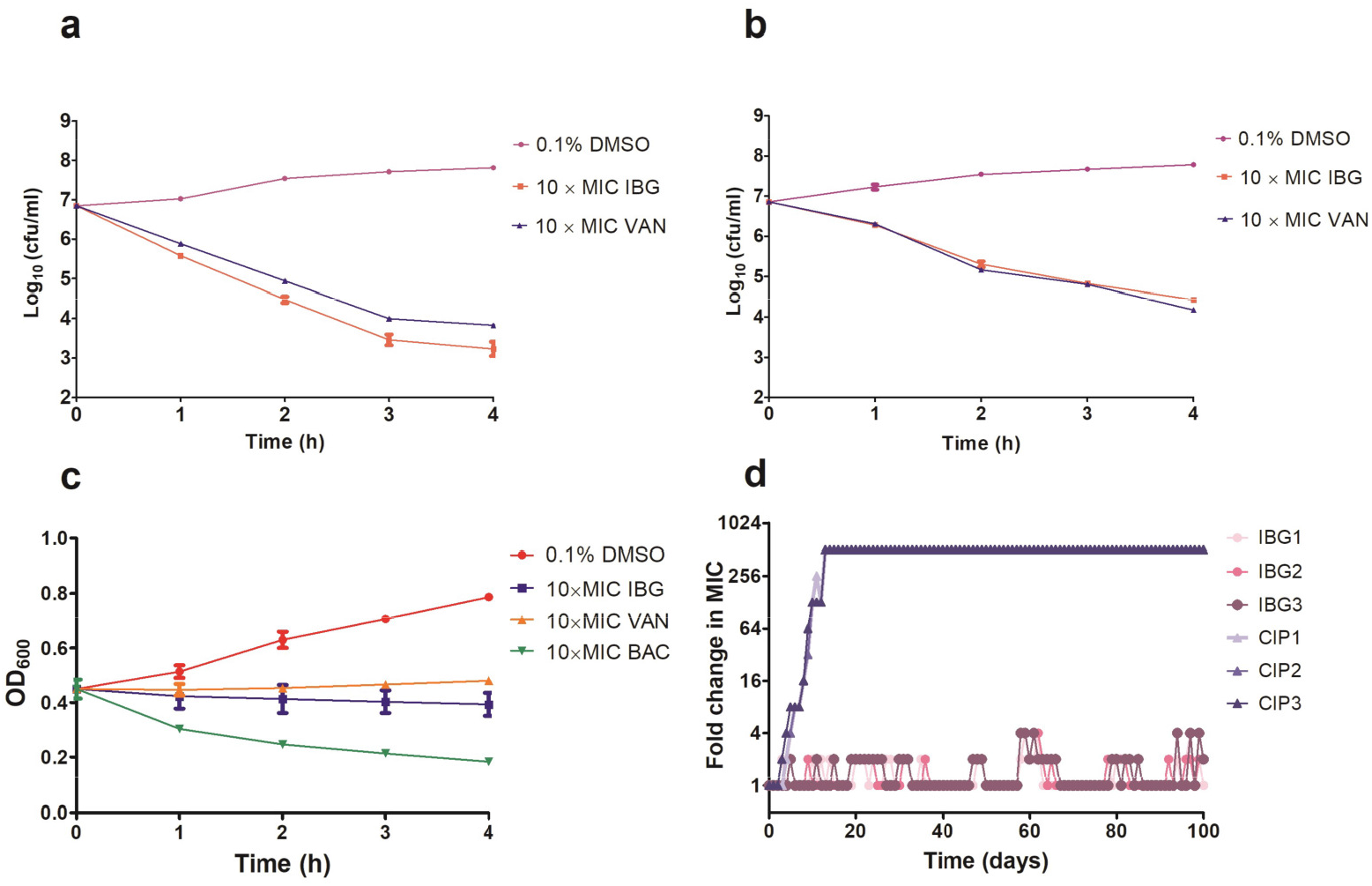
IBG inhibits *S. aureus* growth without detectable mutant development a, Viability of *S. aureus* ATCC 29213 treated with IBG or VAN. b, Viability of MRSA ATCC 43300 treated with IBG or VAN for 4h. c, IBG treatment could not result in *S. aureus* ATCC 29213 lysis. d, Appearance of spontaneous IBG and CIP-resistant *S. aureus* mutants over 100 days of serial passages. IBG, isopropoxy benzene guanidine; VAN, vancomycin; CIP, ciprofloxacin; Individual data points (n = 3 biologically independent experiments) and mean ± s.d. are shown.

### IBG had a low probability of resistance selection in *S. aureus*

We were unable to obtain IBG-resistant mutants by plating 10^10^ CFU of *S. aureus* ATCC 29213 on agar containing 2.5×, 5× or 10× MIC of IBG. Similarly, serial passage of three independent *S. aureus* 29213 cultures (SP1, SP2 and SP3) for 100 days treated with IBG yielded only putative mutants with two or four-fold greater resistance to IBG, whereas serial passage in ciprofloxacin for 100 days generated strains that were 512-fold more resistant (Fig. 2d and Table 3). A total of 24 mutation genes were identified, of which seven encoded hypothetical protein. The mutation strains contained mutations in the genes *atpE*, *glpK*, *isdG-2*, *mnhG1*, *pcrA*, *rsmG*, *rpoE*, *walR*, *lip2* which encode products related to catalytic activity, binding and proton transmembrane transporter activity (Supplementary Table 2).

**Table 3.**
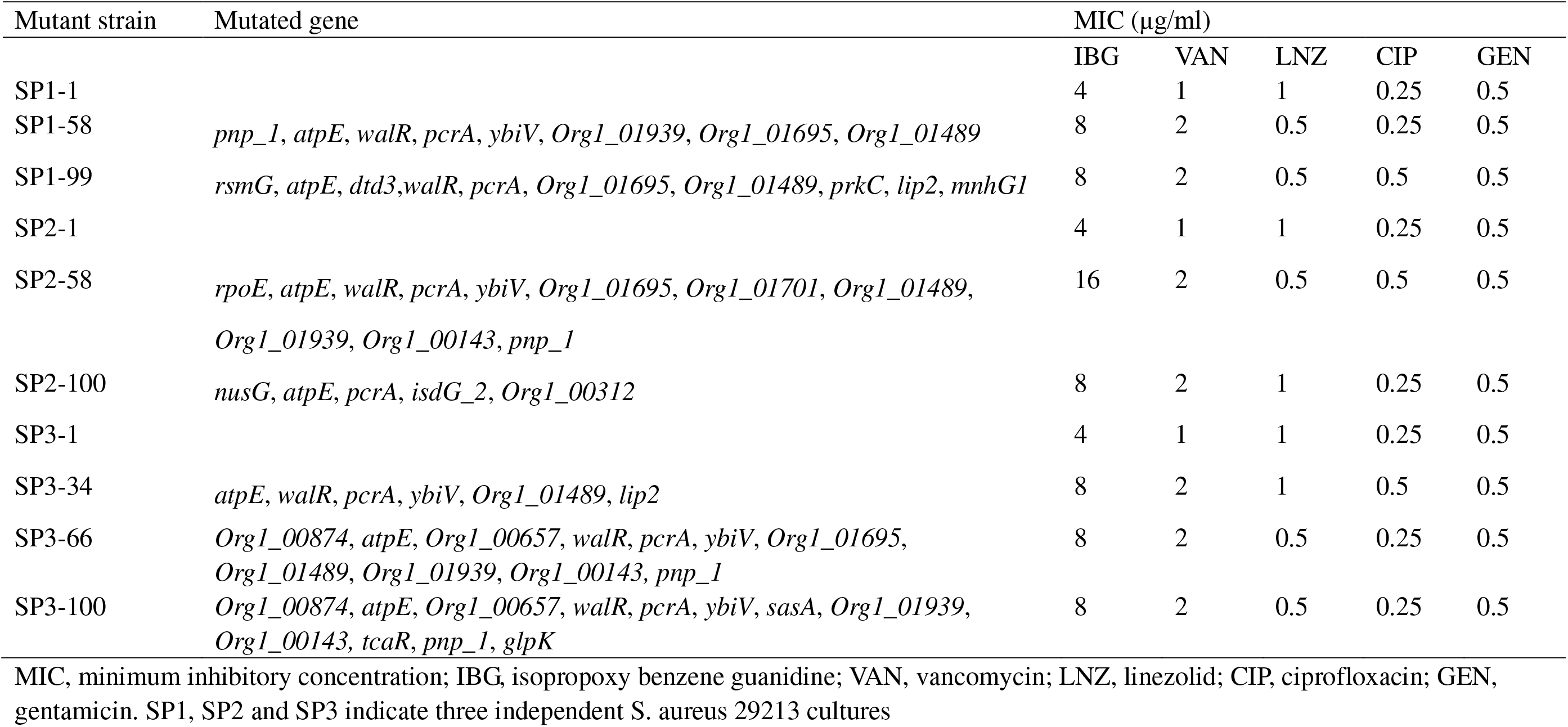
Antimicrobial susceptibility of *S. aureus* ATCC 29213 mutants isolated by serial passage for 100 days

### IBG disrupted cell membrane in multiple ways

DiOC_2_(3) probe has been used to measure the magnitude and stability of ΔΨ in bacterial cells and proteoliposomes. A large reduction in the magnitude of the generated membrane potential was observed in the IBG-treated group, compared to that of the untreated cells and cells in the presence of ampicillin (Fig. 3a). Then we further investigate the membrane disruption properties of IBG by monitoring the SYTOX Green uptake. Our results showed that IBG could clearly permeabilize the bacterial inner membrane even at a concentration of 1× MIC (Fig. 3b). These results were consistent with the findings of TEM. Untreated cells (1% DMSO control) showed a normal cell shape with intact cell walls and membranes. However, remarkable morphological changes in *S. aureus* cells were observed when treated with IBG. Some mesosome-like structures were found in the cytoplasm, cells with ruptured wall and cytoplasmic membrane were also observed. In addition, the cytoplasmic contents of some cells were released to the extracellular medium (Fig. 3c).

**Fig. 3.**
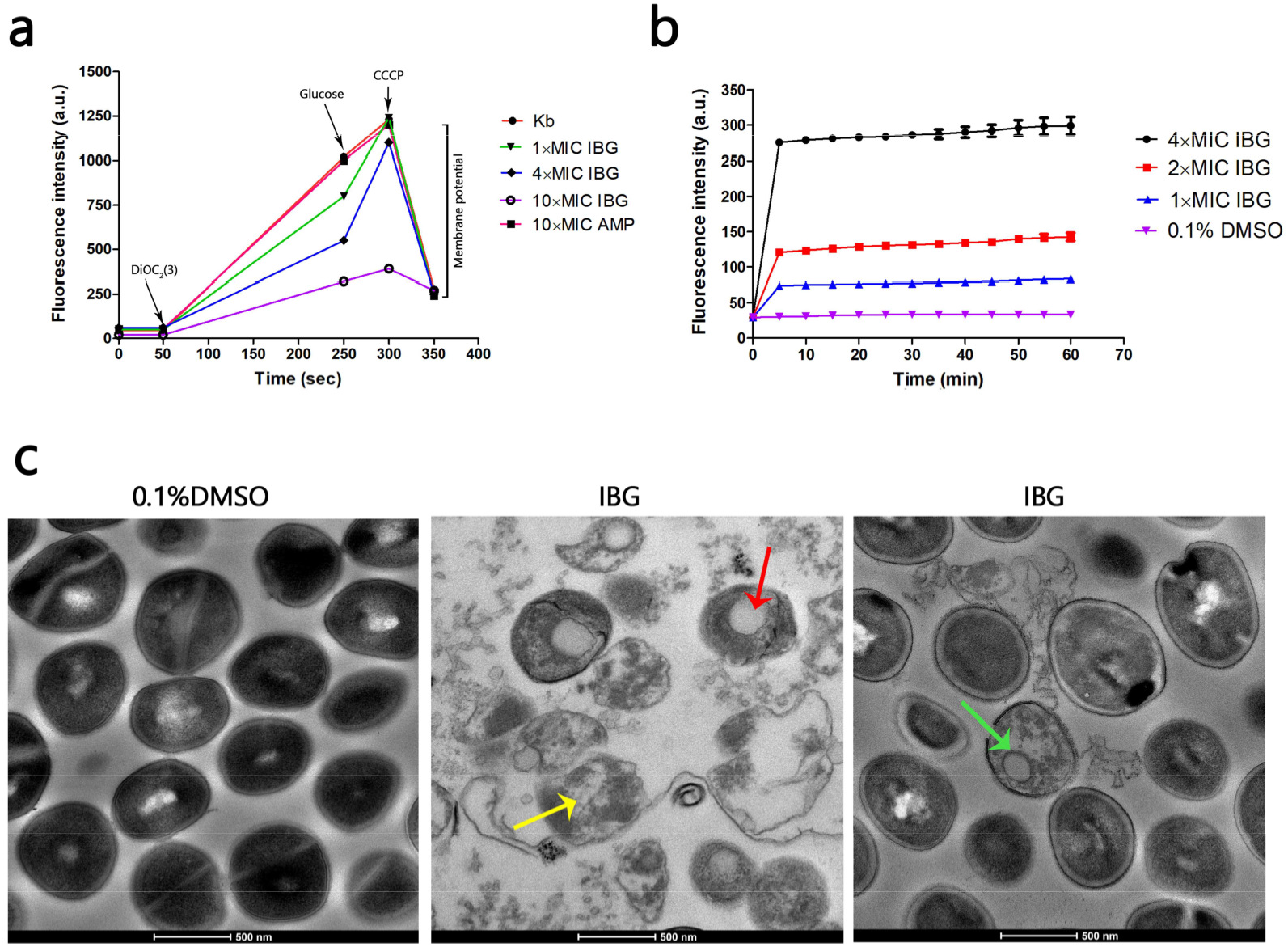
IBG exerts its antibacterial action on the cell membrane of *S. aureus*. a. IBG dissipates the membrane potential of *S. aureus* ATCC 29213. b, Uptake of SYTOX Green by exponential-phase *S. aureus* ATCC 29213 cells treated with IBG. c, Transmission electron micrographs of *S. aureus* cells in response to IBG. Scale bars, 500 nm. These arrows indicate damage to the cell membrane by IBG.

**Fig. 4.**
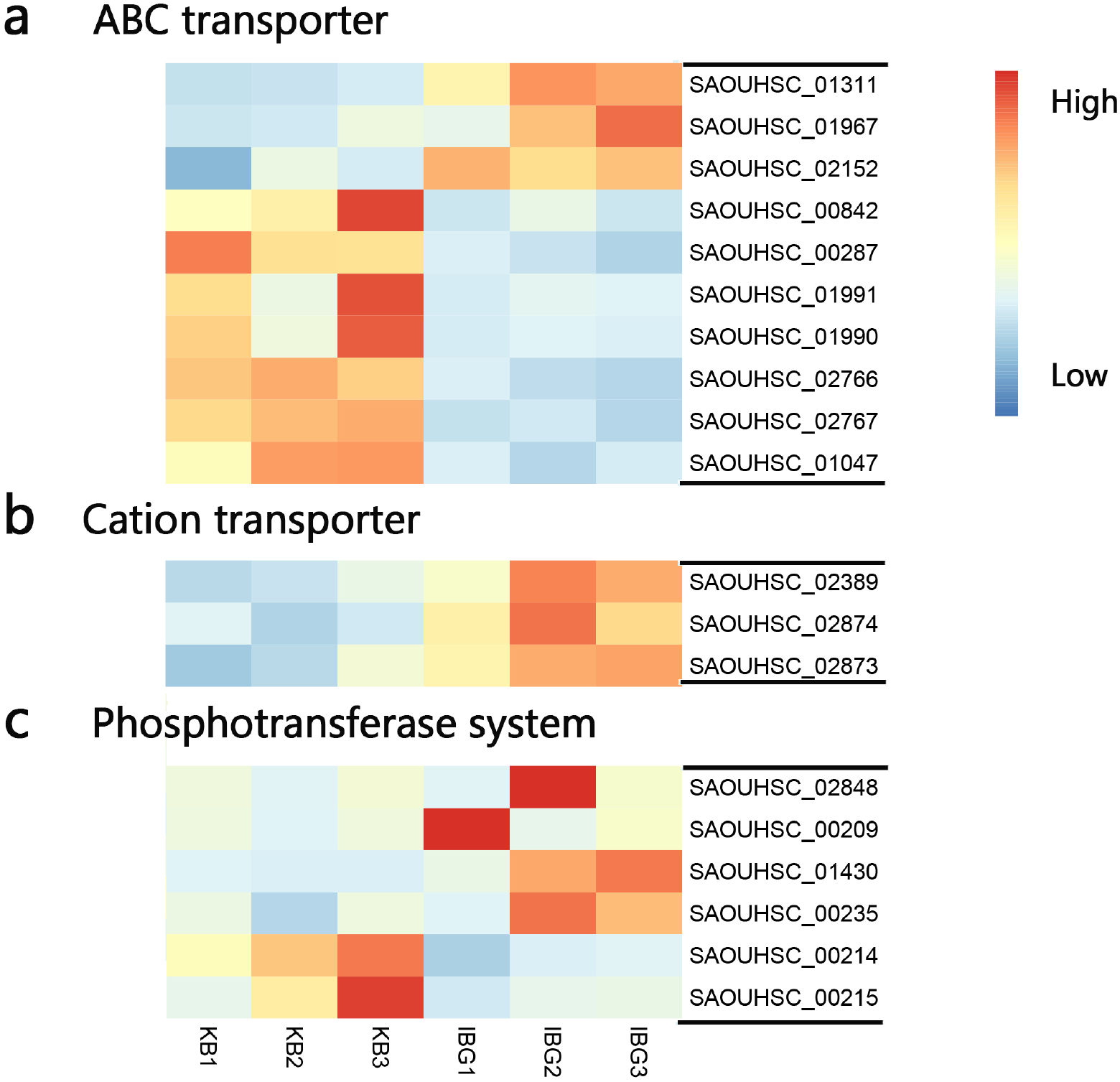
Heat map representation of DGEs involved in cell membrane permeability. a, DGEs related to ABC transpoters. b, DGEs involved in cation transpoters. c, DEGs related to phosphotransferase system. All genes represented have a >2-fold change in expression treated with IBG relative to their expression in control samples.

### Global analysis of transcriptomic response to IBG treatment

The principal component analysis (PCA) analysis revealed that IBG successfully separated treated samples from untreated samples. Compared to the control group, IBG treatment led to 444 DGEs in *S. aureus* ATCC 29213, of which 230 were upregulated and 214 were downregulated (Supplementary Fig. 1).

According to the GO enrichment analysis, the DEGs were involved in diverse processes. As for molecular functions, the main groups that DEGs distributed were catalytic activity, binding and transporter activity. With cellular composition, the top three categories that DEGs located were cell part, membrane and membrane part. While for biological processes, DEGs were mostly involved in metabolic process, cellular process and localization (Supplementary Fig. 5a). In accordance with GO enrichment analysis, subcellular localization results suggested that the majority of DEGs were found in cytoplasm (179 DEGs) and cell membrane (159 DEGs) locations (Supplementary Fig. 5b).

KEGG enrichment analysis indicated that DEGs were mainly involved in pathways of purine metabolism, pyrimidine metabolism, amino sugar and nucleotide sugar metabolism, alanine, aspartate and glutamate metabolism (Fig. 5c). Moreover, according to clueGo analysis, DEGs regulated by IBG treatment were these related to purine metabolism, ABC transporters, pyrimidine metabolism, *S. aureus* infection and alanine, aspartate and glutamate metabolism (Fig. 5d).

**Fig. 5.**
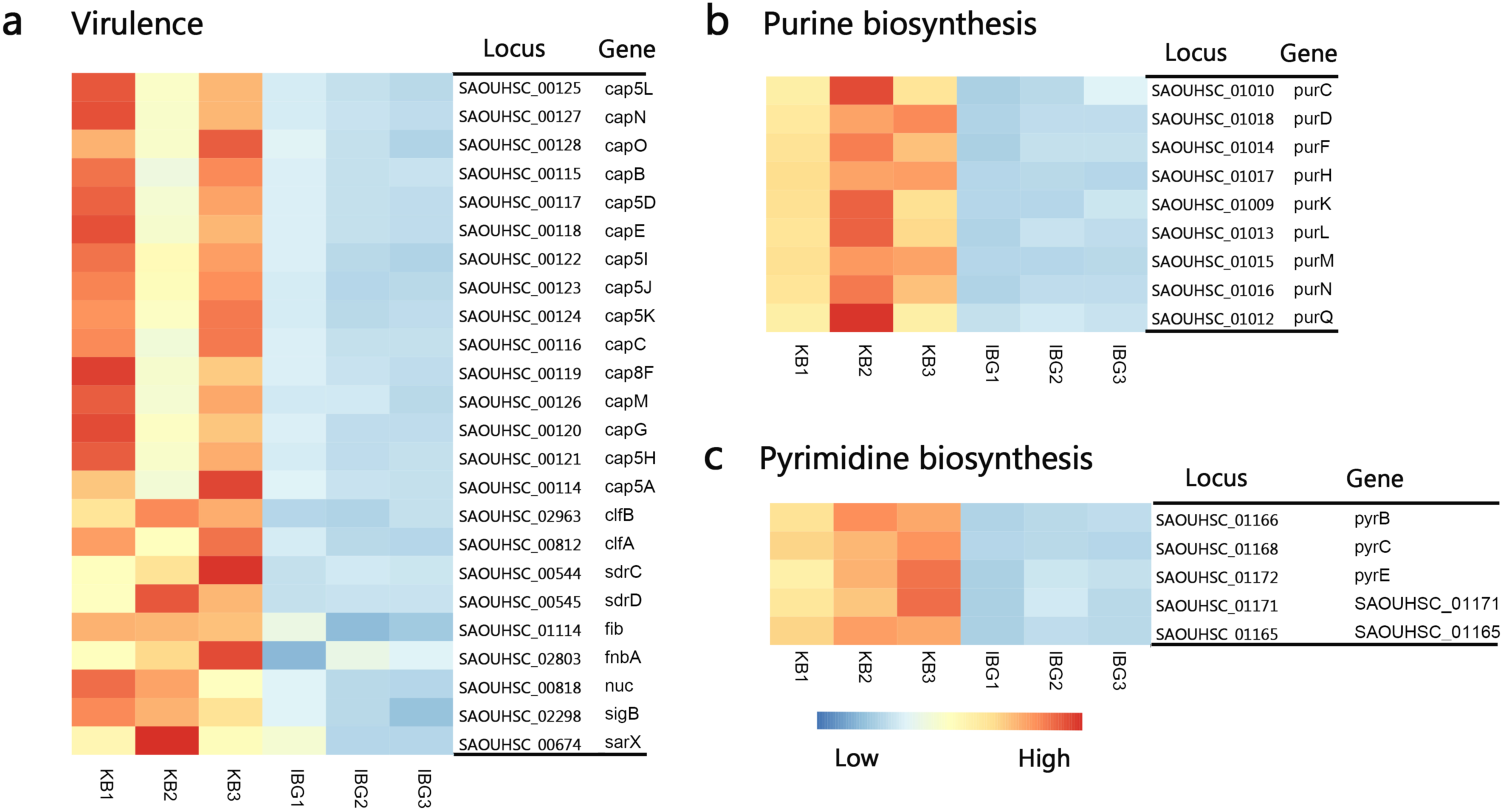
Heat map representation of down-regulated DGEs in response to IBG stress. a, DEGs related to virulence. b, DGEs involved in purine biosynthesis pathway. c, DGEs involved in pyridine biosynthesis pathway. All genes represented have a >2-fold change in expression treated with IBG relative to their expression in control samples.

### Gene related to cell Membrane permeability

According to bioinformatics analysis, a third of DEGs (159/444) were located on cell membrane and most of them coded proteins with membrane transport related function. Functions of these DEGs suggested that most genes were associated with osmotolerance, including those encoding efflux pumps, enzymes involved in ABC transporters, cation transporters and phosphotransferase system (PTS), (Fig. 6).

**Fig. 6.**
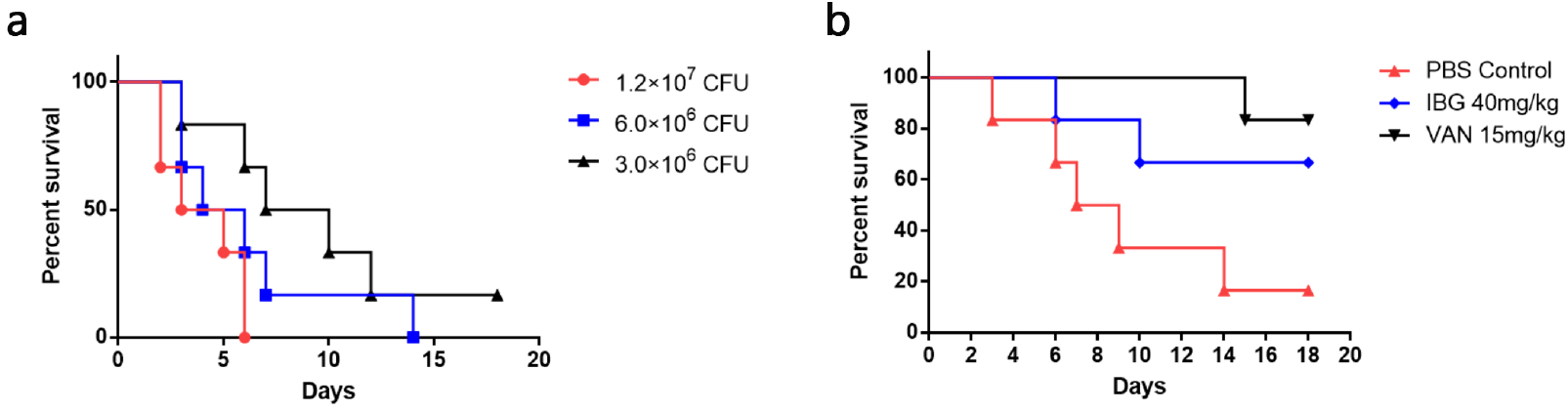
IBG is effective in a mouse model of MRSA septicemic infection. a, Different MRSA infection doses in mouse septicemic model b,Survival curves showing efficacy in a mouse septicemic infection model with the use of IBG.

### IBG down-regulated genes mostly involved in virulence, purine and pyrimidine biosynthesis

The expression of genes involved the virulence of *S. aureus*, including global transcription regulators (staphylococcal accessory regulator *sarX*, RNA polymerase sigma factor *sigB*), cell surface factors (*clfAB*), capsular polysaccharides (*cap5BDEHIJKLNO* and *cap8CGFM*), thermonuclease (*nuc*), cell adhesins (*sdrCD*) was significantly lower in IBG-treated group compared to control. In addition, genes involved in purine biosynthesis pathway (*purC*, *purD*, *purF*, *purH*, *purK*, *purL*, *purM*, *purN*, *purQ*) and pyrimidine biosynthesis pathway (pyrB, pyrC, pyrE, pyrP) were downregulated under IBG treatment (Fig. 7).

### IBG rescues mice from MRSA septicemic infection

The *in vivo* treatment efficacy of IBG was investigated in a mouse septicemia model. Mice were infected with MRSA at a dose ((3.0×10^6^CFU.per mouse)) that leads to 84% of death (Fig. 8a). IBG was introduced at intraperitoneally at 40mg/kg b.w., VAN at a dose of 15mg/kg b.w., or the PBS as a control. The survival rate of infected mice was 66.6% and 83.3% with the dose of IBG or VAN, respectively (Fig. 8). These results suggest that IBG had a potent in protecting mice from septicemic MRSA infection.

### IBG would be used as combination therapy with gentamicin or amikacin

When tested against *S. aureus* ATCC 29213, MRSA ATCC 43300, the FICI ranges were all between 0.5 and 0.75, which means IBG exhibited addition activity when combined with gentamicin, amikacin (Fig. 9).

## Discussion

*Staphylococcus aureus* infections continue to pose a significant challenge to public health due to the diminishing arsenal of effective antibiotics available. Furthermore, the development of novel antibacterial discovery utilizing the traditional approach has not kept pace with the rapid emergence of bacterial resistance to numerous antibiotics, including agents of last resorts. This leads researchers to explore alternative methods to discover new treatment options for bacterial infections; one method that is less time-consuming and more financially viable is repurposing drugs (initially approved for other clinical indications) that possess potent antimicrobial activity. In this study, we showed that isopropoxy benzene guanidine (IBG) displays potent bactericidal activity against *Staphylococcus aureus* by targeting cell membrane.

The successful repurposing of anticoccidial agent robenidine to discover the robenidine analogues as Gram-positive antibacterial agents paved the way for researchers to explore other clinical applications for substituted benzene guanidine derivatives(17). Recent studies, including the present work demonstrate that IBG possesses potent antibacterial activity against an array of important Gram-positive pathogens, including MRSA and multi-drug resistant *Enterococcus*. Additionally, IBG maintained their activity against MRSA isolates exhibiting resistance to ampicillin, penicillin, cefoxitin, florfenicol, erythromycin, tetracycline, valnemulin, ciprofloxacin; this indicates cross-resistance between IBG and these antibiotics is unlikely to occur. It is noted that the antibacterial activity of IBG increased after 24h exposure of the bacteria to the non-ionic detergent (including Triton X-100 and Tris) and ATPase inhibitor DCCD. These findings showed that the antibacterial activity of IBG against *S. aureus* is strengthened by changing the permeability of the cell membrane or with the action of multidrug-resistant (MDR) pumps. Similar phenomenon was also reported by the natural compound sophoraflavanone-B on MRSA (33).

Several reports have indicated that robenidine analogues lack antibacterial activity against Gram-negative bacteria was a consequence of lack of compound penetration of the bacterial outer membrane(17). In this study, we found that IBG was ineffective against Gram-negative bacteria, but showed sensitivity to *E. coli* protoplasts lacking of outer membrane permeability barrier. In addition to this, the sharp reduction in IBG’s MIC was observed in *E. coli* k12 with a deficient efflux pump AcrAB (Supplementary Table 1). Thus the target of IBG is both present in Gram-positive and -negative bacteria and the lack of direct antibacterial activity against Gram-negative bacteria appears to be a byproduct of the barrier imposed by the outer membrane in addition to the presence of active efflux pumps.

Confirmation of IBG’s potent antibacterial activity led us to next explore the potential mechanism of action against *S. aureus*. We attempted to generate a *S. aureus* mutant that is resistant to IBG, but failed. So we performed the serial exposure of bacteria to IBG for 100 days to assess the propensity of bacteria to develop drug resistance. Serial passage of the strain ATCC 29213 treated in ciprofloxacin generated mutant that was 512-fold more resistant than the starting culture (MIC from 0.25 to 128 μg/ml) after only 13 passages, whereas with IBG, the putative mutants were only two or four-fold of its parent strains (MIC from 4 to 16 μg/ml). These results demonstrated that IBG had major advantages over conventional antibiotics and *S. aureus* cannot easily develop resistance to IBG.

Then we performed whole-genome sequences of the resistant mutants (SP1-58, −99, SP2-58,−100, SP3-34, −66, −100) compared to their parental strain. Only the mutations common to all isolates were considered to be associated with resistance. Hence, we identified only two mutations (*aptE* and *pcrA*) shared by all these independently selected resistant isolates. *atpE* gene encodes subunit C of the ATP synthase which utilizes energy stored in the transmembrane electrochemical gradient to synthesize ATP and is important for growth and survival in *S. aureus* (34). The subunit C of bacterial ATP synthase has developed as a clinically validated antimicrobial target and the mutations in *atpE* have been reported to be associated with antibiotic resistance (35). PcrA is a superfamily I DNA helicase which is involved in DNA repair and plasmid rolling circle replication (36). Helicases could translocate along nucleic acids and unwind DNA in reactions that involve the binding and hydrolysis of ATP. PcrA is an ATP-driven 3′-5′ helicase which binds to the 3′ end of DNA and translocates towards the duplex region, opening the dsDNA (37). Strikingly, six of the seven isolates with reduced susceptibility to IBG had single mutations within the *walR* locus. Mutations in *walR* locus have been shown to decrease susceptibility to vancomycin, which is consistent with our result that all the strains exhibited decreased susceptibility to vancomycin after the *walR* mutation. WalR is a transcriptional regulatory protein and a member of the two-component regulatory system WalKR that regulates genes involved in autolysis, biofilm formation and cell wall metabolism. The WalKR system is known to play an important role in resistance to daptomycin, and vancomycin by causing the typical cell wall thickening in *S. aureus* (38). In addition, there is mutation in *rpoE* in the day 34 of SP2. RpoE (δ factor) is the DNA-dependent RNA polymerase (RNAP) subunit presented in certain Gram-positive species and *Firmicutes*. In *S. aureus*, RpoE is part of transcription machinery and most likely interacts directly with the β and/or β′ subunit of RNAP. *RpoE* can be involved in orchestrating the ability of *S. aureus* to react and adapt to environmental changes and thus plays a critical role during virulence (39).

In our previous study, we have confirmed that IBG display potent bactericidal activity against *Enterococcus* by disrupting the cell membrane potential. Hence, we employed a number of techniques to investigate whether IBG has the membrane-bacterial interaction on *S. aureus*. Membrane disruption mediated by IBG is confirmed by the SYTOX green uptake assay, measurement of membrane potential and TEM images. Similar findings were reported in a previous study in which MRSA strains were treated with some compounds or techniques, such as cold-pressed Valencia orange essential oil or ultrasound (26, 40). Moreover, the micrographs indicated that the ultra structural changes caused by IBG treatment might be irreversible. Since an intact cytoplasmic membrane is required for many essential processes in bacteria, perturbation of its structure by IBG maybe the crucial reason for its lethal action on *S. aureus*. However, death is not accompanied by a decline in culture absorbance.

Rencently, transcriptome analysis could simultaneously and globally examine the complete responses at a transcriptional level and have been widely used to explore the antibacterial mechanism of numerous antibiotics. Notably, gene expressions found in other studies were most from low (sublethal) concentrations of antimicrobial agents. Gene expression in bacteria following lethal antimicrobial treatment is hard to find in the published literature (41). In our study, a comparative transcriptomic analysis of *S. aureus* exposure to lethal concentration of IBG showed that 444 genes were differentially expressed, and the major groups to which differentially regulated genes were clustered belong to ABC transport. These contained amino acid ABC transporter ATP-binding protein, ABC transporter ATP-binding protein, peptide ABC transporter permease, exconuclease ABC, spermidine/putrescine ABC transporter permease. ABC transporter systems are associated with nutrient uptake and the export of toxins and antibiotics, and they also have a potential pathogenic role during host infection (42). Previous study has proved that peptide ABC transporter permease, excinuclease ABC and spermidine/putrescine ABC transporter permease are pathogenesis-related ABC transporters and important for *Salmonella* in vivo infection. In addition, PTS transporter subunits were found to be downregulated in IBG treatment. Downregulation of PTS genes is related to decreased cell growth and the transcript levels of PTS component-encoding genes implicated in the utilisation of various sugars under different environmental stresses. Taking these into consideration, it was confirmed that IBG treatment led to increasing membrane permeability in *S. aureus* compared to the control.

In addition, IBG treatment also repressed virulence and ATP-consuming pathways. For instance, purine and pyrimidine biosynthesis pathway genes, crucial for cell growth via DNA and RNA synthesis (43), were downregulated in IBG-treated group. This was probably induced by the derived effects of increased membrane permeability. the down-regulation of virulence factor genes might be very likely as a secondary effect for bacteria to survive. In *S. aureus*, the sigma factor, SigB, represents a powerful regulator to environmental stress, especially in response to antibiotics, and to impact the expression of multiple virulence genes and global regulators, including *sarA* (44). Capsules polysaccharides are the major virulence factors in the pathogenesis of *S. aureus* infections and enable *S. aureus* to avoid swallowing and killing. More than 80% of *S. aureus* isolates express capsular polysaccharide (cap5 or 8) locating in *cap5* or *cap8* gene clusters. In our study, the *cap* genes were all belong to *cap5* or *cap8* gene clusters. Fibronectin-binding protein A (FnbA) and FnbB are known as important adhesions for *S. aureus* and contribute to mediate cellular invasion, and promote virulence during infection (45, 46). Indeed, previous work has demonstrated that the global regulators of *staphylococcal* virulence Agr and Sar coordinate FnbA expression (47). Most interestingly, cold shock protein coding genes *csp* was also markedly repressed. It was confirmed that Csp proteins are involved in survival and the virulence of pathogen.

Its membrane-targeting actions toward bacteria might contribute to the lack of drug resistance during prolonged laboratory culture in its presence because it is difficult for bacteria to remodel their membranes while they are alive. We further moved to confirm the potential of IBG as a therapeutic agent using mice MRSA systemic infection model. These results indicated that IBG had therapeutic potential as a novel antimicrobial agent for drug-resistant Gram-positive bacterial infections. Furthermore, IBG demonstrated additive activity when combined with antibiotics traditionally used to treat systemic MRSA infections. This is important given the emergence of resistance to systemic antimicrobials currently used in the clinic; pairing these antibiotics with IBG may improve the morbidity associated with bacterial infections and stymie the rate at which resistance to these antibiotics arises.

## Conclusions

The study identified a novel compound IBG that possesses potent antimicrobial activity against clinically relevant isolates of *S. aureus*. *In vitro* membrane assays and transcriptome showed that the target of IBG was cell membrane. IBG had a a low probability of resistance selection and considerable efficacy in a mouse septicemic model of MRSA infection. In the summary, IBG has the promising potential to become a new class of antimicrobials for the treatmentof Gram-positive bacterial infections.

## Author contributions

W-G X and Z-L Z conceived this study and designed the experiments. X-F Z and X-P P drafted the manuscript. W-G X and Z-L Z revised the manuscript. X-F P and Z-H Q worked on the synthesis of compound IBG. Y-X L, J H proformed the antimicrobial susceptibility tests, time-kill assay and checkerboard assay. X-F Z, F-P L carried out the RAN-Seq and data analysis. All authors read and approved the final manuscript.

## Reference

1. Årdal C, Balasegaram M, Laxminarayan R, McAdams D, Outterson K, Rex JH, Sumpradit N. 2019. Antibiotic development -economic, regulatory and societal challenges. Nature Reviews Microbiology 2. http://dx.doi.org/10.1038/s41579-019-0293-3.

2. Tacconelli E, Carrara E, Savoldi A, Harbarth S, Mendelson M, Monnet DL, Pulcini C, Kahlmeter G, Kluytmans J, Carmeli Y, Ouellette M, Outterson K, Patel J, Cavaleri M, Cox EM, Houchens CR, Grayson ML, Hansen P, Singh N, Theuretzbacher U, Magrini N, Aboderin AO, Al-Abri SS, Awang JN, Benzonana N, Bhattacharya S, Brink AJ, Burkert FR, Cars O, Cornaglia G, Dyar OJ, Friedrich AW, Gales AC, Gandra S, Giske CG, Goff D, Goossens H, Gottlieb T, Guzman BM, Hryniewicz W, Kattula D, Jinks T, Kanj SS, Kerr L, Kieny MP, Kim YS, Kozlov RS, Labarca J, Laxminarayan R, Leder K, Leibovici L, Levy-Hara G, Littman J, Malhotra-Kumar S, Manchanda V, Moja L, Ndoye B, Pan A, Paterson DL, Paul M, Qiu HB, Ramon-Pardo P, Rodríguez-Baño J, Sanguinetti M, Sengupta S, Sharland M, Si-Mehand M, Silver LL, Song W, Steinbakk M, Thomsen J, Thwaites GE, van der Meer JWM, Van Kinh N, Vega S, Villegas MV, Wechsler-Fördös A, Wertheim HFL, Wesangula E, Woodford N, Yilmaz FO, Zorzet A. 2018. Discovery, research, and development of new antibiotics the WHO priority list of antibiotic-resistant bacteria and tuberculosis. The Lancet Infectious Diseases 18:318–27. http://dx.doi.org/10.1016/s1473-3099(17)30753-3.

3. Turner NA, Sharma-Kuinkel BK, Maskarinec SA, Eichenberger EM, Shah PP, Carugati M, Holland TL, Fowler VGJ. 2019. Methicillin-resistant Staphylococcus aureus an overview of basic and clinical research. Nat Rev Microbiol 17:203–18. http://dx.doi.org/10.1038/s41579-018-0147-4.

4. Nambiar S, Laessig K, Toerner J, Farley J, Cox E. 2014. Antibacterial drug development challenges, recent developments, and future considerations. Clin Pharmacol Ther 96:147–9.. http://dx.doi.org/10.1038/clpt.2014.116.

5. Chin W, Zhong G, Pu Q, Yang C, Lou W, De Sessions PE, Periaswamy B, Lee A, Liang ZC, Ding X, Gao S, Chu CW, Bianco S, Bao C, Tong YW, Fan W, Wu M, Hedrick JL, Yang YY. 2018. A macromolecular approach to eradicate multidrug resistant bacterial infections while mitigating drug resistance onset. Nat Commun 9: 917. http://dx.doi.org/10.1038/s41467-018-03325-6.

6. Trusts TPC. 2019. Antibiotics currently in global clinical development https://www.pewtrustsorg/en/research-and-analysis/data-visualizations/(2014)/antibiotics-currently-in-clinical-development.

7. Ardal C, Baraldi E, Ciabuschi F, Outterson K, Rex JH, Piddock LJV, Findlay D. 2017. To the G20: incentivising antibacterial research and development. The Lancet Infectious diseases 17:799–801. http://dx.doi.org/10.1016/s1473-3099(17)30404-8.

8. Austin BA, Gadhia AD. 2017. New therapeutic uses for existing drugs. Advances in experimental medicine and biology 1031:233–47. http://dx.doi.org/10.1007/978-3-319-67144-4_14.

9. Farha MA, Brown ED. 2019. Drug repurposing for antimicrobial discovery. Nat Microbiol 4:565–77. http://dx.doi.org/10.1038/s41564-019-0357-1.

10. Savoia D. 2016. New antimicrobial approaches reuse of old drugs. Current drug targets 17:731–8.

11. Talbot GH, Jezek A, Murray BE, Jones RN, Ebright RH, Nau GJ, Rodvold KA, Newland JG, Boucher HW. 2019. The Infectious Diseases Society of America’s 10 × ‘20 Initiative (10 New Systemic Antibacterial Agents US Food and Drug Administration Approved by 2020) is 20 × ‘20 a possibility? Clinical infectious diseases an official publication of the Infectious Diseases Society of America 69:1–11. http://dx.doi.org/10.1093/cid/ciz089.

12. Kalia J, Swartz KJ. 2011. Elucidating the molecular basis of action of a classic drug guanidine compounds as inhibitors of voltage-gated potassium channels. Mol Pharmacol 80:1085–95. http://dx.doi.org/10.1124/mol.111.074989.

13. Saczewski F, Balewski L. 2013. Biological activities of guanidine compounds, 2008 - (2012) update. Expert opinion on therapeutic patents 23:965–95. http://dx.doi.org/10.1517/13543776.2013.788645.

14. Flood A, Trujillo C, Sanchez-Sanz G, Kelly B, Muguruza C, Callado LF, Rozas I. 2017. Thiophene/thiazole-benzene replacement on guanidine derivatives targeting alpha2-Adrenoceptors. Eur J Med Chem 138:38–50. http://dx.doi.org/10.1016/j.ejmech.2017.06.008.

15. Massimba-Dibama H, Mourer M, Constant P, Daffe M, Regnouf-de-Vains JB. 2015. Guanidinium compounds with sub-micromolar activities against Mycobacterium tuberculosis. Synthesis, characterization and biological evaluations. Bioorg Med Chem 23:5410–8. http://dx.doi.org/10.1016/j.bmc.2015.07.053.

16. Kantor S, Kennett RL, Jr., Waletzky E, Tomcufcik AS. 1970. 1,3-Bis (p-chlorobenzylideneamino) guanidine hydrochloride (robenzidene) new poultry anticoccidial agent. Science 168:373–4.

17. Abraham RJ, Stevens AJ, Young KA, Russell C, Qvist A, Khazandi M, Wong HS, Abraham S, Ogunniyi AD, Page SW, O’Handley R, McCluskey A, Trott DJ. 2016. Robenidine analogues as gram-positive antibacterial agents. J Med Chem 59:2126–38. http://dx.doi.org/10.1021/acs.jmedchem.5b01797.

18. Ogunniyi AD, Khazandi M, Stevens AJ, Sims SK, Page SW, Garg S, Venter H, Powell A, White K, Petrovski KR, Laven-Law G, Totoli EG, Salgado HR, Pi H, Coombs GW, Shinabarger DL, Turnidge JD, Paton JC, McCluskey A, Trott DJ. 2017. Evaluation of robenidine analog NCL195 as a novel broad-spectrum antibacterial agent. PLoS One 12: e0183457. http://dx.doi.org/10.1371/journal.pone.0183457.

19. Zhang X, Han D, Pei P, Hao J, Lu Y, Wan P, Peng X, Lv W, Xiong WG, Zeng ZL. 2019. In vitro antibacterial activity of isopropoxy benzene guanidine against multidrug-resistant Enterococci. Infect Drug Resist 12:3943–53. http://dx.doi.org/10.2147/IDR.S234509.

20. Li W, Liu JH, Zhang XF, Wang J, Ma ZB, Chen L, Zeng ZL. 2018. Emergence of methicillin-resistant Staphylococcus aureus ST398 in pigs in China. Int J Antimicrob Agents 51:275–6. http://dx.doi.org/10.1016/j.ijantimicag.2017.10.013.

21. 2018. Performance Standards for Antimicrobial Susceptibility Testing, M100. 28th informational supplement. Clinical and Laboratory Standards Institute, Wayne, PA.

22. Mohammad H, Reddy PV, Monteleone D, Mayhoub AS, Cushman M, Seleem MN. 2015. Synthesis and antibacterial evaluation of a novel series of synthetic phenylthiazole compounds against methicillin-resistant Staphylococcus aureus (MRSA). Eur J Med Chem 94:306–16. http://dx.doi.org/10.1016/j.ejmech.2015.03.015.

23. Ling LL, Schneider T, Peoples AJ, Spoering AJ, Engels I, Conlon BP, Mueller A, Schaberle TF, Hughes DE, Epstein S, Jones M, Lazarides L, Steadman VA, Cohen DR, Felix CR, Fetterman KA, Millett WP, Nitti AG, Zullo AM, Chen C, Lewis K. 2015. A new antibiotic kills pathogens without detectable resistance. Nature 517:455–9. http://dx.doi.org/10.1038/nature14098.

24. Kim W, Zhu W, Hendricks GL, Van Tyne D, Steele AD, Keohane CE, Fricke N, Conery AL, Shen S, Pan W, Lee K, Rajamuthiah R, Fuchs BB, Vlahovska PM, Wuest WM, Gilmore MS, Gao H, Ausubel FM, Mylonakis, E. 2018. A new class of synthetic retinoid antibiotics effective against bacterial persisters. Nature 556:103–7. http://dx.doi.org/10.1038/nature26157.

25. Wang Y, Mowla R, Guo L, Ogunniyi AD, Rahman T, De Barros Lopes MA, Ma S, Venter H. 2017. Evaluation of a series of 2-napthamide derivatives as inhibitors of the drug efflux pump AcrB for the reversal of antimicrobial resistance. Bioorganic & Medicinal Chemistry Letters 27:733–9. http://dx.doi.org/10.1016/j.bmcl.2017.01.042.

26. Li J, Ahn J, Liu D, Chen S, Ye X, Ding T. 2016. Evaluation of ultrasound-induced damage to Escherichia coli and Staphylococcus aureus by flow cytometry and transmission electron microscopy. Applied and environmental microbiology 82:1828–37. http://dx.doi.org/10.1128/aem.03080-15.

27. Chou K-C, Shen H-B. 2010. Cell-PLoc 2.0 an improved package of web-servers for predicting subcellular localization of proteins in various organisms. Natural Science 2:1090–103. http://dx.doi.org/10.4236/ns.2010.210136.

28. Bindea G, Mlecnik B, Hackl H, Charoentong P, Tosolini M, Kirilovsky A, Fridman WH, Pages F, Trajanoski Z, Galon J. 2009. ClueGO: a Cytoscape plug-in to decipher functionally grouped gene ontology and pathway annotation networks. Bioinformatics (Oxford, England) 25:1091–3. http://dx.doi.org/10.1093/bioinformatics/btp101.

29. Thomas VM, Brown RM, Ashcraft DS, Pankey GA. 2019. Synergistic effect between nisin and polymyxin B against pandrug-resistant and extensively drug-resistant Acinetobacter baumannii. Int J Antimicrob Agents. http://dx.doi.org/10.1016/j.ijantimicag.2019.03.009.

30. Falagas ME, Voulgaris GL, Tryfinopoulou K, Giakkoupi P, Kyriakidou M, Vatopoulos A, Coates A, Hu Y. 2019. Synergistic activity of colistin with azidothymidine against colistin-resistant Klebsiella pneumoniae clinical isolates collected from inpatients in Greek hospitals. Int J Antimicrob Agents. http://dx.doi.org/10.1016/j.ijantimicag.2019.02.021.

31. Thangamani S, Mohammad H, Abushahba MF, Sobreira TJ, Hedrick VE, Paul LN, Seleem MN. 2016. Antibacterial activity and mechanism of action of auranofin against multi-drug resistant bacterial pathogens. Sci Rep 6: 22571. http://dx.doi.org/10.1038/srep22571.

32. Abdelraouf K, Chavda KD, Satlin MJ, Jenkins SG, Kreiswirth BN, Nicolau DP. 2020. Piperacillin-tazobactam-resistant/third-generation Cephalosporin-susceptible Escherichia coli and Klebsiella pneumoniae isolates Resistance Mechanisms and In vitro-In vivo Discordance. Int J Antimicrob Agents 105885. http://dx.doi.org/10.1016/j.ijantimicag.2020.105885.

33. Mun SH, Joung DK, Kim SB, Park SJ, Seo YS, Gong R, Choi JG, Shin DW, Rho JR, Kang OH, Kwon DY. 2014. The mechanism of antimicrobial activity of sophoraflavanone B against methicillin-resistant Staphylococcus aureus. Foodborne pathogens and disease 11:234–9. http://dx.doi.org/10.1089/fpd.2013.1627.

34. Balemans W, Vranckx L, Lounis N, Pop O, Guillemont J, Vergauwen K, Mol S, Gilissen R, Motte M, Lancois D, De Bolle M, Bonroy K, Lill H, Andries K, Bald D, Koul A. 2012. Novel antibiotics targeting respiratory ATP synthesis in Gram-positive pathogenic bacteria. Antimicrob Agents Chemother 56:4131–9. http://dx.doi.org/10.1128/aac.00273-12.

35. Lamontagne Boulet M, Isabelle C, Guay I, Brouillette E, Langlois JP, Jacques PE, Rodrigue S, Brzezinski R, Beauregard PB, Bouarab K, Boyapelly K, Boudreault PL, Marsault E, Malouin F. 2018. Tomatidine is a lead antibiotic molecule that targets Staphylococcus aureus ATP synthase subunit C. Antimicrob Agents Chemother 62:0066–4804. http://dx.doi.org/10.1128/aac.02197-17.

36. Dillingham MS, Soultanas P, Wiley P, Webb MR, Wigley DB. 2001. Defining the roles of individual residues in the single-stranded DNA binding site of PcrA helicase. Proc Natl Acad Sci U S A 98:8381–7. http://dx.doi.org/10.1073/pnas.131009598.

37. Mhashal AR, Choudhury CK, Roy S. 2016. Probing the ATP-induced conformational flexibility of the PcrA helicase protein using molecular dynamics simulation. Journal of molecular modeling 22: 54. http://dx.doi.org/10.1007/s00894-016-2922-3.

38. Howden BP, McEvoy CR, Allen DL, Chua K, Gao W, Harrison PE, Bell J, Coombs G, Bennett-Wood V, Porter JL, Robins-Browne R, Davies JK, Seemann T, Stinear T. P.2011. Evolution of multidrug resistance during Staphylococcus aureus infection involves mutation of the essential two component regulator WalKR. PLoS pathogens 7: e1002359. http://dx.doi.org/10.1371/journal.ppat.1002359.

39. Weiss A, Ibarra JA, Paoletti J, Carroll RK, Shaw LN. 2014. The delta subunit of RNA polymerase guides promoter selectivity and virulence in Staphylococcus aureus. Infection and immunity 82:1424–35. http://dx.doi.org/10.1128/IAI.01508-14.

40. Muthaiyan A, Martin EM, Natesan S, Crandall PG, Wilkinson BJ, Ricke SC. 2012. Antimicrobial effect and mode of action of terpeneless cold-pressed Valencia orange essential oil on methicillin-resistant Staphylococcus aureus. Journal of applied microbiology 112:1020–33. http://dx.doi.org/10.1111/j.1365-2672.2012.05270.x.

41. Wu S, Yu PL, Wheeler D, Flint S. 2018. Transcriptomic study on persistence and survival of Listeria monocytogenes following lethal treatment with nisin. J Glob Antimicrob Resist 15:25–31. http://dx.doi.org/10.1016/j.jgar.2018.06.003.

42. Liu M, Feng M, Yang K, Cao Y, Zhang J, Xu J, Hernandez SH, Wei X, Fan M. 2020. Transcriptomic and metabolomic analyses reveal antibacterial mechanism of astringent persimmon tannin against Methicillin-resistant Staphylococcus aureus isolated from pork. Food chemistry 309: 125692. http://dx.doi.org/10.1016/j.foodchem.2019.125692.

43. Fleury B, Kelley WL, Lew D, Gotz F, Proctor RA, Vaudaux P. 2009. Transcriptomic and metabolic responses of Staphylococcus aureus exposed to supra-physiological temperatures. BMC Microbiol 9: 76. http://dx.doi.org/10.1186/1471-2180-9-76.

44. Tuchscherr L, Bischoff M, Lattar SM, Noto Llana M, Pfortner H, Niemann S, Geraci J, Van de Vyver H, Fraunholz MJ, Cheung AL, Herrmann M, Volker U, Sordelli DO, Peters G, Loffler B. 2015. Sigma Factor SigB Is Crucial to Mediate Staphylococcus aureus Adaptation during Chronic Infections. PLoS pathogens 11: e1004870. http://dx.doi.org/10.1371/journal.ppat.1004870.

45. Shinji H, Yosizawa Y, Tajima A, Iwase T, Sugimoto S, Seki K, Mizunoe Y. 2011. Role of fibronectin-binding proteins A and B in in vitro cellular infections and in vivo septic infections by Staphylococcus aureus. Infection and immunity 79:2215–23. http://dx.doi.org/10.1128/iai.00133-11

46. Goncheva MI, Flannagan RS, Sterling BE, Laakso HA, Friedrich NC, Kaiser JC, Watson QW, Wilson CH, Sheldon JR, McGavin MJ, Kiser PK, Heinrichs DE. 2019. Stress-induced inactivation of the Staphylococcus aureus purine biosynthesis repressor leads to hypervirulence. Nat Commun 10: 775. http://dx.doi.org/10.1038/s41467-019-08724-x.

47. Wolz C, Pohlmann-Dietze P, Steinhuber A, Chien YT, Manna A, van Wamel W, Cheung A. 2000. Agr-independent regulation of fibronectin-binding protein(s) by the regulatory locus sar in Staphylococcus aureus. Molecular microbiology 36:230–43. http://dx.doi.org/10.1046/j.1365-2958.2000.01853.x.

